# Comparison of alternative models of human movement and the spread of disease

**DOI:** 10.1101/2019.12.19.882175

**Authors:** Ottar N. Bjørnstad, Bryan T. Grenfell, Cecile Viboud, Aaron A. King

**Affiliations:** Center for Infectious Disease Dynamics, Pennsylvania State University, University Park, Pensylvannia, USA; Department of Ecology and Evolutionary Biology, Princeton University, Princeton, New Jersey, United States of America; Division of International Epidemiology and Population Studies, Fogarty International Center, National Institutes of Health, Bethesda, Maryland, USA; Department of Ecology and Evolutionary Biology, Center for the Study of Complex Systems, and Department of Mathematics, University of Michigan, Ann Arbor, Michigan, USA

## Abstract

Predictive models for the spatial spread of infectious diseases has received much attention in recent years as tools for the management of infectious diseas outbreaks. Prominently, various versions of the so-called gravity model, borrowed from transportation theory, have been used. However, the original literature suggests that the model has some potential misspecifications inasmuch as it fails to capture higher-order interactions among population centers. The fields of economics, geography and network sciences holds alternative formulations for the spatial coupling within and among conurbations. These includes Stouffer’s rank model, Fotheringham’s competing destinations model and the radiation model of Simini et al. Since the spread of infectious disease reflects mobility through the filter of age-specific susceptibility and infectivity and since, moreover, disease may alter spatial behavior, it is essential to confront with epidemiological data on spread. To study their relative merit we, accordingly, fit variants of these models to the uniquely detailed dataset of prevaccination measles in the 954 cities and towns of England and Wales over the years 1944-65 and compare them using a consistent likelihood framework. We find that while the gravity model is a reasonable first approximation, both Stouffer’s rank model, an extended version of the radiation model and the Fotheringham competing destinations model provide significantly better fits, Stouffer’s model being the best. Through a new method of spatially disaggregated likelihoods we identify areas of relatively poorer fit, and show that it is indeed in densely-populated conurbations that higher order spatial interactions are most important. Our main conclusion is that it is premature to narrow in on a single class of models for predicting spatial spread of infectious disease. The supplemental materials contain all code for reproducing the results and applying the methods to other data sets.

**Author summary:** The ability to predict how infectious disease will spread is of great importance in the face of the numerous emergent and re-emergent pathogens that currently threatening human well-being. We identified a variety of alternative models that predict human mobility as as a function of population distribution across a landscape. These consider some models that account for pair-wise interactions between population centers, as well as some that allow for higher-order interactions. We trained the models using a uniquely rich spatiotemporal data set on pre-vaccination measles in England and wales (1944-65), which comprises more than a million records from 954 cities and towns. Likelihood rankings of the different models reveal strong evidence for higher-order interactions in the form of competition among cities as destinations for travelers and, thus, dilution of spatial transmission. The currently most commonly used so-called ‘gravity’ models were far from the best in capturing spatial disease dynamics.

## Introduction

Accurately predicting the geographical spread of emerging, re-emerging, and recurrent epidemics in an increasingly globalized world is a matter of international urgency in the wake of outbreaks of emerging pathogens such as severe acute respiratory syndrome (SARS), Middle East respiratory syndrome (MERS), and ebola virus disease (EBVD) as well as re-emerging and resurgent pathogens such as influenza subtype A-H1N1, whooping cough, and measles. In response to this challenge, various ‘spatial interaction’ models describing human movement as a function of population distribution have been proposed. Some of these borrow from economics and human geography while others adapt models from movement ecology and the physics of reaction and diffusion on heterogenous landscapes.

In recent years, the field has seen the widespread adoption of a family of so-called ‘gravity’ models from transportation theory and human geography (see, e.g., [1, 2]). In its most common form, the gravity model posits that the migration flux between a pair of cities is log-linearly dependent on their respective sizes and on their separating distance. The application of this simple model to disease spread was originally proposed by Murray and Cliff [3], but over the last decade, many studies have used it to explain historical, or predict future, disease spread in a range of infections including measles [4], influenza [5–7], cholera [8], and yellow fever [9]. While its application has yielded insights, the geography literature has highlighted a prominent shortcoming of the gravity model. In particular, these models ignore the potential for competitive or synergistic interactions among population centers (Fig. 1; [10, 11]). Thus, for example, movement between Boston and Washington DC is assumed to be unaffected by the presence of the intervening city of New York.

**Fig 1.**
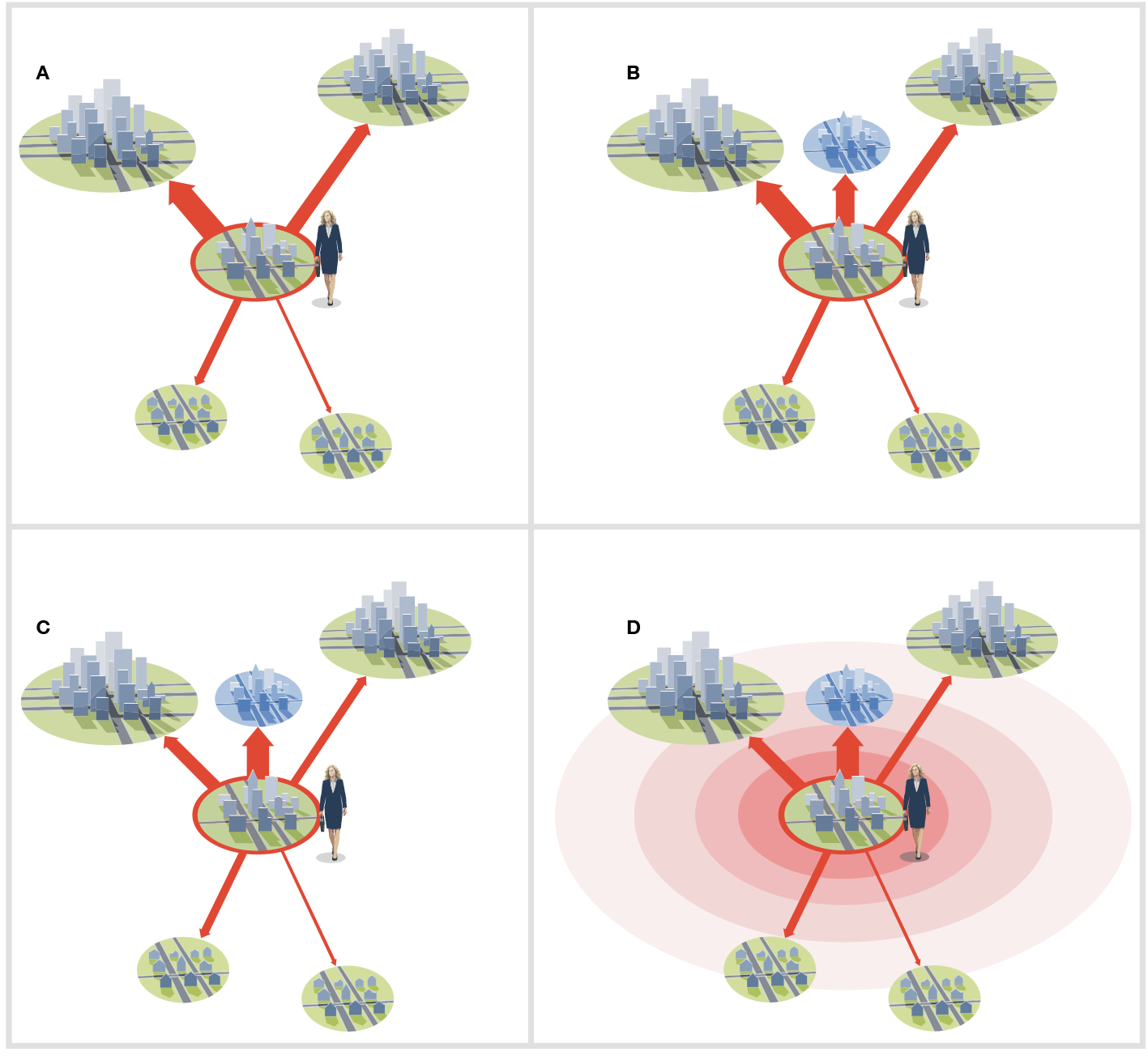
**Spatial interaction models** predict the flux of human movements between population centers (cities, towns, villages) as a function of the distribution of the population on the landscape. In this diagram, the relative magnitude of the fluxes from a focal town to other population centers are represented by the widths of the arrows. In the widely-employed gravity models (A), interactions among cities is strictly pairwise. Thus the addition of a new town (B) has no effect on the movement to other towns. In Fotheringham’s competing destinations model (C), however, competition or synergy among nearby communities can reduce or augment fluxes. Stouffer’s model of intervening opportunities and the radiation model (D) posit that movement from one city to another is diminished by the presence of opportunities in communities more proximal to the source city.

As it happens, there are distinct families of models, arising from economics and geography, that predict human movement patterns while allowing for higher-order interactions among cities. In particular, Stouffer’s [12] ‘law of intervening opportunities’ posits that “the number of persons going a given distance is directly proportional to the number of opportunities at that distance and inversely proportional to the number of intervening opportunities”. This idea has given rise to alternative models for human movement patterns [13] including a recent incarnation as the ‘radiation’ model [14].

While these models have proved useful in their original applications, their use in describing infectious disease spread is complicated by two considerations. First, spatial interaction models predict bulk migration fluxes between population centers, but the impact of these movements on the dynamics of infectious disease can depend strongly not only on the magnitude, but also on the composition, of the migrant pool. In particular, the extent to which migrants are more or less likely to be susceptible to infection, or actually infected, than the general population can be critically important, as can the age profile of the migrant pool, due to pronounced age-specific patterns of behavior, susceptibility, and infectiousness [5, 7, 15]. Second, infectious disease dynamics can feed back onto migration. Most obviously, disease symptoms can influence movement behavior, as seen, for example, in the fact that a mild cold may induce minimal changes in movement behavior but severe hemorrhagic fever or acute paralytic disease will typically slow the movement of the infected hosts. These two considerations do not preclude the utility of spatial interaction models in the infectious disease context; they do, however, complicate their use and the importance of understanding *transmission relevant* migration fluxes, which will be some sort of ‘effective average’ of movement as filtered through such aforementioned complications.

Several spatial interaction models have been parameterized and tested using various mobility data such as commuter flows (e.g., [5, 14]), mobile phone geolocations (e.g., [16]), social media (e.g., [13]), and microsimulations (e.g., [17]). However, in view of the challenges just noted, the ultimate test of the models is against data on the actual spread of infection rather than bulk movement of people or cell phones. Bjørnstad and Grenfell [18] proposed that for acute immunizing infections, the spatio-temporal patterns of fade-outs (i.e., local disease extinction) across metapopulations provide valuable information on disease spread because these patterns reflect spatial transmission unclouded by local transmission. In this paper, we extract information from fade-out patterns and a consistent likelihood framework to compare and contrast to a suite of models including (i) the gravity model [1], (ii) Fotheringham’s competing destinations model [10], (iii) Stouffer’s rank model [12], and (iv) the radiation model [14].. We confront these models weekly data on measles incidence from all 954 cities and towns in England and Wales from 1944 to 1965 [4, 19]. Comparing fits and predictions, we show that while the gravity model is a reasonable first approximation, Stouffer’s rank model, an extended version of the radiation model, and the competing destinations model all provide significantly better fits, Stouffer’s model performing the best.

## Materials and Methods

### Data

Historical incidence of measles in England and Wales has been an influential testbed for models and methods in disease dynamics since Bartlett’s [20, 21] seminal work on its recurrent epidemics. We use the spatially resolved weekly measles data across all 954 cities and towns of England and Wales from 1944, when notification was made mandatory by the UK Registrar General (OPCS), until 1965, which saw subtle shifts in political boundaries around London. Vaccination was not introduced in the UK until 1967, so that these data span a period where measles dynamics were unaffected by mass immunization. The data set is complete except for a region-wide underreporting rate of around 50% [22, 23]. Grenfell et al. [19] give a detailed description of the data; the entire data set has been made available by Lau et al. [24].

An important feature of the system is that, between 29% and 38% of the population (c. 47M during this period) resided in a small number (13–28 depending on exact definition) of communities above a critical community size (CCS) of c. 250–300k. Cities larger than the CCS tend to sustain local chains of transmission. The remaining 60–70% were distributed among the more than 900 communities smaller than the CCS where local extinctions are more or less frequent (depending on population size and degree of isolation) and, consequently, the rate of reintroduction of the pathogen via spatial transmission is an important determinant of measles incidence. In ecological terms, the prevaccination measles system represented an epidemic mainland-island metapopulation (e.g., [25, 26]). Our analysis exploits this fact, using the timing and spatial pattern and timing of reintroductions to inform the parameters of each of the spatial interaction models.

### Local dynamics

Using a spatially-extended time series susceptible-infected-recovered (TSIR) framework (e.g., [4, 27]), we can calculate, for each population center, the probability that a spatial interaction happens (i.e., a contact between a resident susceptible host and a non-resident infectious host) *and* that the contact results in a new local chain of transmission [18]. In previous analyses, we showed that the signature of such events will be drowned out in the presence of endemic circulation (see [23]). Following a local extinction, however, the re-colonization rate contains critical information on the spatial interactions.

The spatially-extended TSIR model predicts that the expected number of new infected hosts in community *i* in epidemic generation *t* + 1 will be

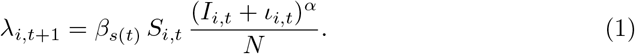

Here, *β*_*s*(*t*)_ is the seasonal transmission rate as molded by opening and closing of schools and the proportion of transmission that occurs within the school setting as the school year progresses [23]; *s*(*t*) = t mod 26 is the seasonality function; *S_i,t_/N* is the probability that a local individual is susceptible; *I_i,t_* is the local number of infections; *α* is a correction for the discrete-time approximation of the underlying continuous-time process [28]; and *ι_i,t_* is the sum of transmission-relevant spatial interactions i has with the other communities in the epidemic metapopulation. Following a local extinction, *I_i,t_* = 0, so this expectation is

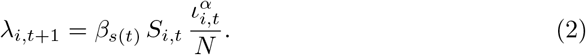

Assuming demographic stochasticity in transmission due to the underlying epidemic birth-and-death process, the realized number of cases in the next generation will be [23]:

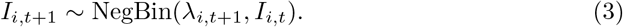

As a consequence, the number of susceptibles in the next generation will be

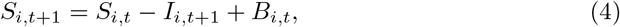

where *B_i,t_* is the recruitment of local susceptibles through births during the generation interval. The ‘serial interval’ for measles is 10–14 days [29], so we follow previous TSIR analyses in aggregating the weekly data in 2-week increments (though [23] investigate shorter intervals). Using susceptible reconstruction methods [30], we estimate all local parameters using the method of Finkenstadt et al. [31]. The online supplement contains full documentation of these analyses and code for exact replication of all results presented in this paper.

Conditioning on the local parameters and restricting analyses at each location to biweeks in which measles incidence was zero, we use Eqs. 2 and 4 to form a negative binomial likelihood to estimate all parameters of each of the candidate spatial interaction models.

### Spatial interaction models

When including special cases, such as pure diffusion and variants, we consider a total of 10 spatial interaction models (Fig. 1). Each of these amounts to a specification of the *ι* term in Eqs. 1 and 2.

#### Gravity model

Under the gravity model, the spatial interaction between locations *i* and *j* takes the form 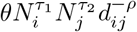, where *θ, τ*_1_, *τ*_2_, and *ρ* are non-negative parameters. This leads to the following formulation for disease-relevant spatial interactions:

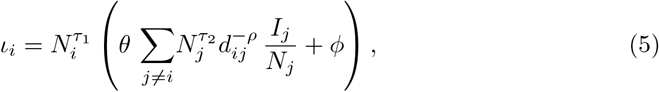

where the sum is across all non-self potential donor communities, and *I_j_/N_j_* is the fraction of infected individuals in donor community *j*. The gravity model has two important special cases: *ρ* = 0, *τ*_1_ = *τ*_2_ = 1, which is a mean-field model, and *τ*_1_ = *τ*_2_ = 0, which is simple spatial diffusion. The *ϕ* parameter represents background spatial transmission that is not predictable on the basis of distance and size [24].

#### Xia’s model

In the original analysis of the spatiotemporal dynamics of measles, Xia et al. [4] used a formulation rooted in the gravity-model literature but with a slightly different formulation:

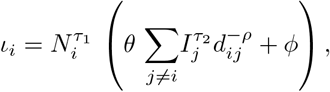

which is the same as Eq. 5 if *τ*_2_ = 1, but not otherwise.

#### Competing destinations model

Fotheringham [11] noted that gravity models may be misspecified because they only consider pairwise interactions among locations. He argued that nearby or intervening destinations may, in fact, make spatial interactions either more or less likely. A synergistic effect occurs, for example, if individuals are disproportionately inclined to do all their shopping in districts with many shops. Antagonistic effects may arise where travel between two cities, say Boston and Washington, is made less likely by the presence of an intervening city such as New York because individuals from either Boston or Washington can get their out-of-town needs fulfilled by visiting New York. Under Fotheringham’s [11] competing destinations formulation the flux between *i* and *j* takes the form 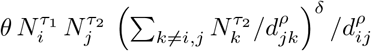. As in the gravity model, *τ*_1_ and *τ*_2_ control how ‘eagerness to travel’ and city ‘attractiveness’ scales with population size; and *ρ* measures how the likelihood of travel decays with distance. The parameter *δ* quantifies how destinations *k*, of various sizes, at distances *d_jk_* from the donor *j*, modulate the spatial interaction between recipient *i* and and donor *j*. In particular, *δ* > 0 indicates a synergistic effect; *δ* < 0, an antagonistic effect. The resultant formulation for disease-relevant spatial interactions between community *i* and everywhere else is:

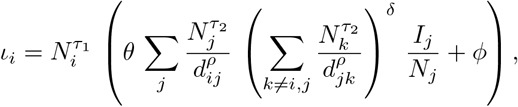

where *ϕ* again represents background, spatially unpredictable spread.

#### Stouffer’s rank model

Noulas et al. [13] recently proposed that human mobility patterns should be studied using Stouffer’s [12] rank (“law of intervening opportunities”) model. Stouffer argued against absolute distance as a direct influence on spatial interactions. Rather, he argued the decline of coupling with distance emerges due to the accumulation of intervening opportunities as the distance between two locations increases. To apply this notion in our measles metapopulation context, we use population size as a proxy for “opportunities”. Accordingly, we let Ω(*i, j*) be the collection of towns located closer to town *i* than is town *j*: Ω(*i, j*) = {*k*: 0 < *d*(*i, k*) ≤ *d*(*i, j*)}. The Stouffer model for disease-relevant spatial interactions is, then

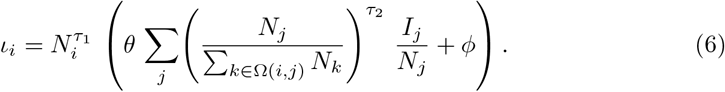

Stouffer’s original model was framed in the context of continuous distribution of population across a landscape. In the present metapopulation context, we therefore have a choice as to whether the opportunities offered by location *i* itself should be counted among the intervening opportunities. In Eq. 6, it is excluded. If instead we allow the set Ω(*i, j*) to include *i* – that is, within-community opportunities reduce spatial coupling – we arrive at what we shall term the ‘Stouffer variant’ model: Ω(*i, j*) = {*k*: 0 ≤ *d*(*i, k*) ≤ *d*(*i, j*)}

#### Extended radiation model

The radiation model proposed by Simini et al. [14] was derived independently but is in spirit related to the original ideas of Stouffer [12]. Our metapopulation version of this model with respect to disease-relevant spatial interactions is

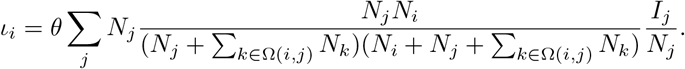

As with the Stouffer model, we consider the set Ω(*i, j*) to either exclude (‘radiation”) or include (‘radiation variant”) community size *i* itself in the spatial interaction formulation. As originally formulated [14], these models have but a single free parameter for the spatially-structured component, the overall spatial interaction strength, *θ*; To facilitate comparison with other models, and since component of spatially random spread has been implicated in measles metapopulation dynamics [24], we also entertain a third variant of the radiation model (‘extended radiation’) that includes a spatially-random background rate (*ϕ*):

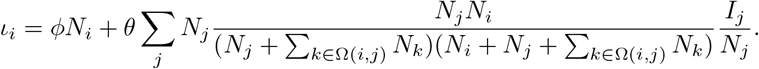

Of the models considered, the gravity variants (including mean-field and pure diffusion) include purely pairwise interactions whilst the other models allow for higher-order interactions either explicitly (as in the competing destination model) or implicitly (as in the Stouffer and radiation models).

### Statistical inference

We use maximum likelihood to estimate the parameters of each spatial interaction model and profile likelihoods to study correlation among parameters and possible identifiability issues [27]. For model comparison we use quasi-likelihoods [32] to accommodate over-dispersion;and rank models according to their quasi Akaike Information criterion (QAIC) scores [33]. We calculate the quasi-likelihood variance-inflation-factor (*ĉ* in Table 1) from the Pearson goodness-of-fit statistic according to the recommendations of McCullagh and Nelder [32].

**Table 1.** MLE estimates. Models: SV – Stouffer variant, XR – extended radiation, CD – competing destinations, S – Stouffer, G – gravity, X – Xia, RV – radiation variant, R – radiation, MF – mean field, D – diffusion. Here, *ℓ* denotes the log likelihood, c is the computed variance inflation factor, ΔQAIC is the QAIC relative to that of the best-fitting model, and the other parameters are as in the model equations.

| Model | $\ell$ | $\hat{c}$ | $\Delta\text{QAIC}$ | $\log_{10} \theta$ | $\log_{10} \phi$ | $\rho$ | $\tau_1$ | $\tau_2$ | $\delta$ | $\psi$ |
| --- | --- | --- | --- | --- | --- | --- | --- | --- | --- | --- |
| SV | -196466.8 | 2.85 | 0 | -0.80 | -4.6 |  | 0.88 | 1.73 |  | 5.07 |
| XR | -196630.9 | 3.19 | 111.0 | -1.07 | -5.1 |  |  |  |  | 5.10 |
| CD | -196654.3 | 2.89 | 135.5 | 0.44 | -3.7 | 1.86 | 0.66 | 0.50 | -1.01 | 5.08 |
| S | -196778.9 | 2.76 | 218.8 | -0.82 | -4.3 |  | 0.82 | 1.44 |  | 5.14 |
| G | -197733.0 | 2.88 | 889.9 | -3.45 | -3.4 | 1.52 | 0.61 | 0.13 |  | 5.27 |
| X | -198028.0 | 2.85 | 1096.8 | -6.02 | -6.3 | 0.82 | 0.61 | 0.43 |  | 5.33 |
| RV | -199682.2 | 3.95 | 2248.8 | -0.82 |  |  |  |  |  | 5.61 |
| R | -200228.5 | 4.37 | 2631.9 | -0.91 |  |  |  |  |  | 5.71 |
| MF | -200369.7 | 4.10 | 2737.0 | -8.77 | -5.1 |  |  |  |  | 5.74 |
| D | -202735.5 | 2.02 | 4394.0 | -0.72 | -0.8 | 1.94 |  |  |  | 6.26 |

As per the recommendation of Burnham and Anderson [33], we use the c of the best-fitting model to adjust the QAICs. The analysis is based on the 274,943 data points that are relevant to the hazard likelihood i.e., all biweeks immediately following a biweek of local disease absence [18]) All calculations are documented in the online supplement, which includes code for the detailed replication of all results.

## Results

Table 1 shows estimated parameters and likelihoods for each of the spatial interaction models. The most obviously interesting result is the ranking of the models in terms of explanatory power. The two variants of the gravity model fall in the middle of the pack; both are significantly inferior to the Stouffer variant, extended radiation and competing destinations models, as judged by the QAICs. The evidence, therefore, suggests strongly that higher-order interactions influence the spatial spread of measles in the prevaccination era. The *δ* parameter of the competing destination model is significantly negative suggesting that the disease transportation network is dominated by antagonistic spatial effects. The diffusion (*τ*_1_ = *τ*_2_ = 0) and mean-field (*ρ* = 0) models are special cases of several of the models, and these are clearly far inferior in accounting for the data. The original single-parameter radiation model variants performed relatively poorly. However, when augmented with an additional parameter to allow for spatially-random background seeding (‘extended radiation’) placed second between Stouffer’s rank model and Fotheringham’s competing destinations model. This may not be too surprising since the radiation model may in spirit be seen as a reincarnation of Stouffer’s original argument.

As discussed by Jandarov et al. [27], there are identifiability issues associated with the parameterizing the various spatial interaction models for infection spread. This is in the from of ridges in the likelihood surfaces over model parameter spaces (see Supplement). Since each model amounts to a parameterization of the 954 × 954 matrix of interaction among England and Wales’ cities and towns, the likelihood ridges mean that different parameter combinations may map on to similar spatial interaction networks. To explore the drivers among model differences in fit, we therefore contrast model predictions of disease import and export rates. Fig. 2 shows the mean predicted export and import rates vs. population size for each of the models. The dependence of mean export rate on population size contrasts somewhat among models (Fig. 2A), but the importation rates follow similar patterns (Fig. 2B). Despite their differences in underlying mathematical equations, there must be more nuanced differences among the models that contribute their relative fit/lack-of-fit. This begs the question: “What are these differences and are they biologically meaningful or spatially random?”

**Fig 2.**
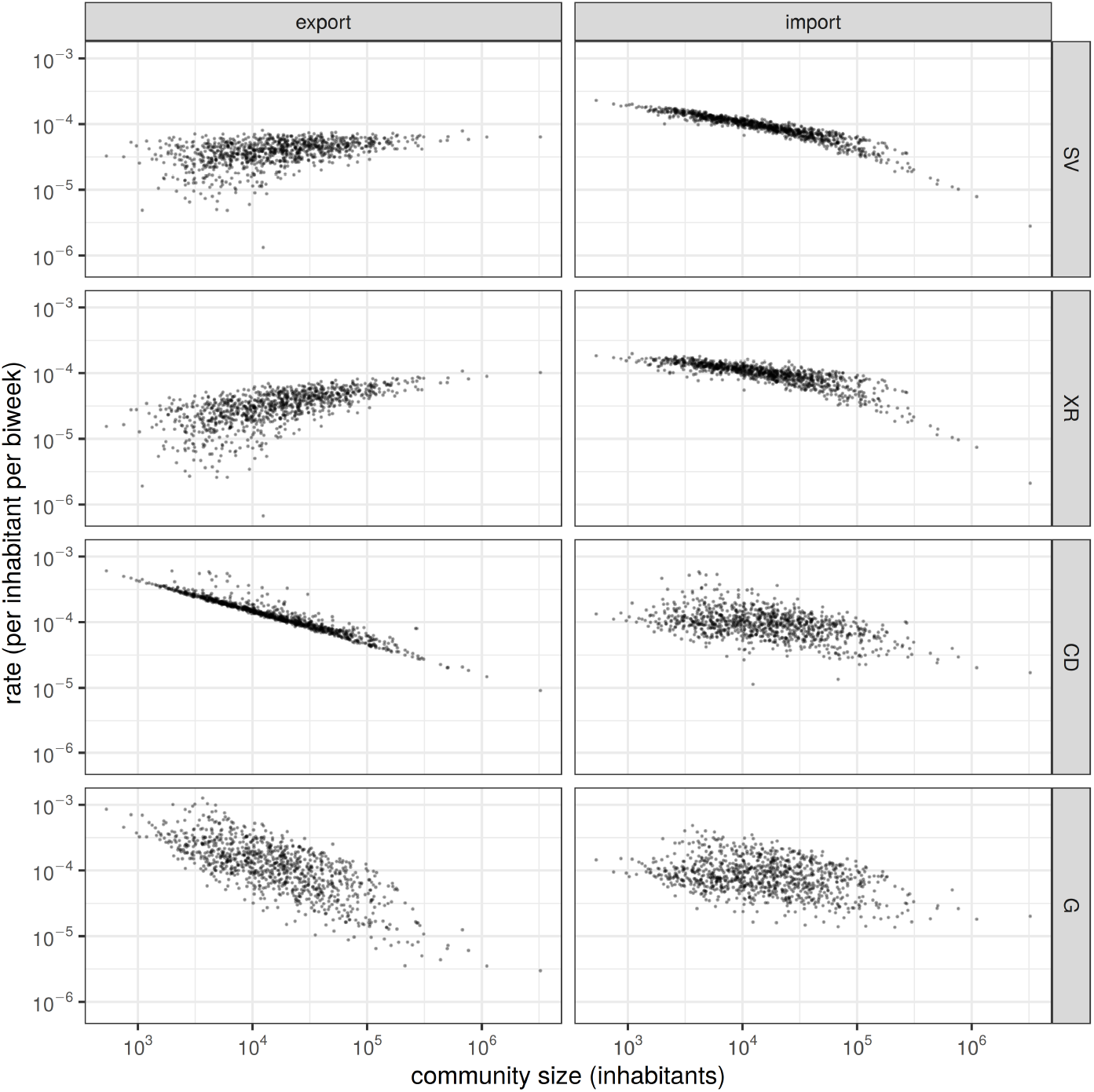
Model-predicted mean import and export rates vs. community size. Models: SV – Stouffer variant, XR – extended radiation, CD – competing destinations, G – gravity.

A cursory review of the recent literature suggests that the gravity model is the most prominent in the infectious disease context, followed by the radiation model. Stouffer’s model and the competing destinations model have rarely been applied in this field. It is therefore useful to use the gravity model fit as a baseline and explore how the better-fitting models diverge from this baseline. Comparing matrices with half a million entries is very difficult, so we amploy a new ‘spatial likelihood contrast’ (SLiC) method with which to study the relative merit of the different spatial interaction models. The idea is to disaggregate the overall hazard likelihoods by individual cities to study how particular communities contribute to improved or diminished relative fit of each model. To do so we normalize each location’s contribution to the overall likelihood by the number of data points each conurbation contributes to the likelihood (the accumulated stretches of measles absence) and then map the model-model differences onto the landscape (Fig. 3). Inspection of Fig. 3 suggests that the poorer fit of the gravity model vs. the Stouffer or competing destinations models is primarily in the urban northwest. The supplement provides the full set of pairwise SLiC contrasts among all models. To test whether the apparent patterns are statistically significant, we compute local indicators of spatial association (LISA; [34]) statistics for all population centers with fewer than 50k inhabitants (Fig. 3A, B). The main failure of the gravity model is in predicting coupling among the cities and towns of the Manchester-Liverpool-Leeds conurbation northwestern England where higher-order antagonistic interactions are evidently an important influence. Comparing predicted export (Fig. 3C) and import (Fig. 3D) rates from the Stouffer vs. gravity model suggest that the significant SLiC statistic is driven by the gravity model’s over-prediction of infection export rates in that region.

**Fig 3.**
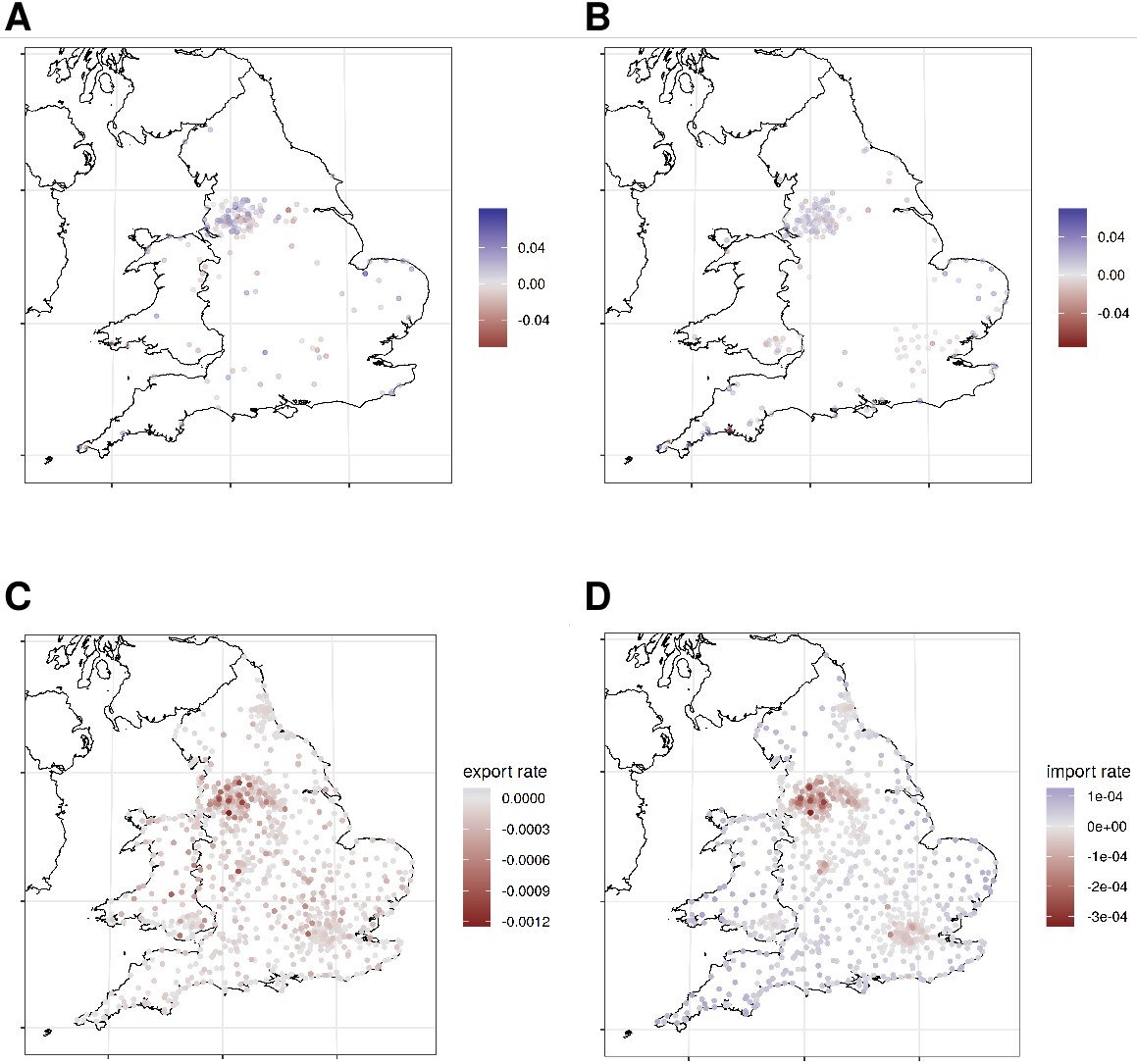
Locations wherein, on average, the (A) Stouffer variant model and (B) competing destinations model outperform the gravity model according to the LISA statistic applied to the spatial likelihood contrasts of all towns of size <50k. Differences in predicted (C) export and (D) import rates of the Stouffer variant vs. gravity model.

## Discussion

Invasion and re-invasion of immunizing human infections often occurs within a metapopulation context because of the aggregation of the human host into more or less discrete population centers. The threat of pandemics and emergent infections has motivated a need to understand and predict patterns of spatial spread of infections among such centers. Several candidate models have been proposed to make such predictions. In wildlife systems—rabies being a classic example—relatively simple diffusion models have performed well in describing the speed of spatial spread (e.g. [35, 36]). In the case of epidemics playing out within a population center, models closer to the mean field model (i.e., with uniform connectivity) often do remarkably well (see [37] for a social network perspective on this). However, such models generally do not capture regional spread of human infections because of the complicated patterns of human movement (the third wave of the 2009 influenza pandemic across the USA, perhaps, being an unusually diffusive counterexample [7]). To elucidate these patterns, many empirical studies have been performed and a variety of abstractions have been proposed. The task of choosing among these abstractions for help in help predicting spread of infectious disease is complicated by the fact that (i) model formulations may differ with respect to how well they actually reproduce aggregate mobility patterns, and (ii) model fit may be shaped by the manner in which mobility is filtered by transmission and behavior.

To advance the discussion of these issues, we considered a suite of candidate models of spatial interactions and fit them to the pre-vaccination measles data from England and Wales using a spatial hazards approach [18]. We considered four main classes of models, including the gravity, competing destinations, Stouffer rank, and radiation models, in addition to pure diffusion and mean field models which are interesting special cases. The gravity model—which has gained recent prominence in infectious disease modeling—assumes that migration flux between pairs of cities depends on their sizes and separating distance, but is unaffected by the presence of other centers. The other three models all allow spatial interactions to be influenced size and proximity of other communities. By challenging these models to emulate patterns in the uniquely rich spatiotemporal data on incidence of measles in English and Welsh cities and towns between 1944 and 1965, we obtain intriguing new insights. Although the gravity model can capture many broad spatiotemporal patterns [4], among the alternative formulations, both Stouffer’s [12] rank model and Fotheringham’s [10] competing destinations model and a background-augmented variant of Simini et al’s [14] radiation model outperformed it. The original radiation model [14] did less well than gravity and simple mean-field and diffusion models worse still, in testimony to the complexity of human mobility in governing the spread of childhood infectious disease.

As the gravity model has had a very rapid adoption in spatial disease epidemiology during the last decade (though Noulas et al. [13] have noted the importance of considering alternatives), we used this as a baseline. Despite its common usage it significantly under-performs relative to the several other models. Combining SLiC and LISA statistics we found that the greatest tension between the gravity model and the preferred alternatives is in the Manchester-Liverpool-Leeds conurbation of northwestern England, where many towns and villages commingle amongst several major cities. While ranked lower than Stouffer’s model, Fotheringham’s competing destinations model reveals two interesting patterns. First, *δ* ≈ −1 is an ndication of competition among nearby towns for travellers. Second, the numerical estimates of *τ*_1_ and *τ*_2_ in this model appear to echo the finding of Lau et al. [24] (who used a related gravity formulation) that spatial interaction, once discounted for higher-order interference, is roughly proportional to the geometric mean of the community sizes. This curious suggestion warrants future investigation.

In this study we have used actual disease incidence data, as opposed to raw measurments of human movement, to further our understanding of the principles governing how human hosts spread ifectious disease across a populated landscape. We have found both strong evidence of higher-order interactions among population centers, and that candidate models differ significantly in their ability to capture the empirical patterns. We hope our findings will help stimulate a systematic, data-driven discussion of the relative merit of alternative predictive models for the probable path for spatial spread of infectious disease. Clearly, the prospects are good for good for further refinements and improved parameterizations.

## Acknowledgments

This study has benefitted from financial support from the National Science Foundation, the National Institutes of Health, the Bill and Melinda Gates Foundation and Fogarty International Center’s program on Research and Policy for Infectious Disease Dynamics (RAPIDD). AAK was supported by Grant #R01AI101155 from the U.S. National Institute of 376 Allergy and Infectious Diseases and by Grant #U54GM111274 from the U.S. National 377 Institute of General Medical Sciences.

## Disclaimer

CV: This paper does not necessarily represent the views of the NIH or the US government.

## Supporting Information Legends

The supporting information is a compiled html Rmarkdown to exactly reproduce all the calculations in the manuscript.

